# An *in vivo* model of echovirus-induced meningitis in neonates

**DOI:** 10.1101/2022.02.11.480094

**Authors:** Alexandra I. Wells, Carolyn B. Coyne

## Abstract

Echoviruses are amongst the most common causes of aseptic meningitis worldwide, which can cause long-term sequelae and death, particularly in neonates. However, the mechanisms by which these viruses induce meningeal inflammation is poorly understood, owing at least in part to the lack of *in vivo* models that recapitulate this aspect of echovirus pathogenesis. Here, we developed an *in vivo* neonatal mouse model that recapitulates key aspects of echovirus-induced meningitis. We found that expression of the human homologue of the neonatal Fc receptor (FcRn), the primary echovirus receptor, in neonatal mice was not sufficient for infection of the brain. However, ablation of type I, but not III, IFN signaling in mice expressing human FcRn permitted high levels of echovirus replication in the brain, with corresponding clinical symptoms including delayed motor skills and hind limb weakness. We also defined the immunological response of the brain to echovirus infections and identified key cytokines induced by this infection. Lastly, we found that echoviruses robustly replicate in the leptomeninges, where they induce profound inflammation and cell death. Together, this work establishes an *in vivo* model of aseptic meningitis associated with echovirus infections and defines the specificity of echoviral infections within the meninges.

**Significance Statement:** Echoviruses are amongst the most common causes of aseptic meningitis worldwide, which can cause long-term sequelae or even death. The mechanisms by which echoviruses infect the brain are poorly understood, largely owing to the lack of robust *in vivo* models that recapitulate this aspect of echovirus pathogenesis. Here, we establish a neonatal mouse model of echovirus-induced aseptic meningitis and show that expression of the human homologue of the neonatal Fc receptor (FcRn), the primary receptor for echoviruses, and ablation of type I interferon (IFN) signaling are required to recapitulate echovirus-induced meningitis and clinical disease. These findings provide key insights into the host factors that control echovirus-induced meningitis and a model that could be used to test anti-echovirus therapeutics.

## Introduction

Enteroviruses are the main causes of aseptic meningitis worldwide, which is characterized by meningeal inflammation not associated with any identifiable bacterial species in cerebrospinal fluid (CSF). Approximately 90% of aseptic meningitis cases in infants (1) and 50% in older children and adults (2) are caused by enteroviruses, with the group B enterovirus members coxsackievirus B (CVB) and echoviruses being amongst the most common (3, 4). Infants and young children are particularly vulnerable to complications of echovirus-associated neuronal complications, which can cause long-term sequelae including seizure disorders (5, 6), and are associated with high rates of mortality, which occurs in as many as one-third of cases (7, 8). Despite the clear association between echoviruses and aseptic meningitis, the mechanisms by which these viruses induce meningeal inflammation is poorly understood, owing at least in part to the lack of *in vivo* models that recapitulate this aspect of echovirus pathogenesis.

Echoviruses are the largest subgroup of the Enterovirus genus and consist of approximately 30 serotypes. We and others have shown that the neonatal Fc receptor (FcRn) is the primary receptor for echoviruses (9, 10). FcRn transports and regulates the circulating half-life of immunoglobulin G (IgG) and albumin and is enriched in the endothelium of the central nervous system (CNS), including the blood-brain barrier (BBB), where it mediates the efflux of IgG from the brain (11-15). Although several studies have investigated the possible mechanistic basis for CVB-associated neuronal dysfunction *in vitro* and *in vivo* (16-18), much less is known about echovirus-associated CNS complications. Intracerebral inoculation of newborn mice expressing VLA-2, a reported receptor for echovirus 1, exhibit paralysis and motor defects (19). Other work suggests that type I interferons (IFNs) play a role in echovirus 11 CNS disease following intracranial inoculation (20). We previously generated an *in vivo* mouse model of echovirus 11 pathogenesis using adult and neonatal mice and showed that expression of the human homologue of FcRn in mice lacking type I IFN signaling were susceptible to echovirus infection, including infection of the brain (21). However, the consequences of echovirus infection including the induction of cytokines and cell death in the brains of these mice, as well as identification of the region(s) of the brain targeted by infection were not explored.

The meninges surround the brain and are composed of three distinct membranous layers which includes the dura, arachnoid, and pia mater. A hallmark of aseptic meningitis involves inflammation of the meninges, resulting in immune cell infiltration and swelling. In human cases of confirmed echovirus aseptic meningitis, infection is associated with a robust inflammatory response, as indicated by the presence of high levels of proinflammatory mediators in CSF (22-24). Higher levels of type I IFNs are also present in CSF isolated from enterovirus-associated meningitis than from bacterial meningitis (25), suggesting that these IFNs play a prominent role in aseptic meningitis. Type I IFNs, which include IFNαs and IFN-β, provide key antiviral defenses from many neurotropic viruses, including flaviviruses, alphaviruses, and herpesviruses (26). In some cases, type III IFNs (IFN-λs) also defend from CNS viral infections (27, 28) and have been proposed to function by alterations in BBB permeability (27). Whether type I and III IFNs play differential roles in echovirus infections of the CNS is unknown.

Here, we developed an *in vivo* model of echovirus-induced aseptic meningitis in neonatal mice which recapitulates many of the disease manifestations observed in humans. We show that expression of human FcRn alone is not sufficient to mediate echovirus infection of the brains of 7-day-old neonatal mice. In addition, the brains of mice deficient in either type I or III IFN signaling alone were not permissive to echovirus infection. In contrast, we found that humanized FcRn mice deficient in IFNAR, the type I IFN receptor, but not IFNLR, the type III IFN receptor, exhibited high levels of echovirus replication in the brain, with corresponding clinical symptoms including hind limb weakness and paralysis. Using this model, we defined the immune response in the brains of echovirus-infected mice, which included the induction of high levels of IL-6, CXCL10, and granulocyte colony-stimulating factor (GSF)-3. Lastly, we show that echoviruses replicate in the leptomeninges and induce inflammation and cytotoxicity in these membranes, including activation of apoptotic cell death. Together, this work establishes an *in vivo* model of aseptic meningitis associated with echovirus infections and defines the specificity of echoviral infections in the meninges.

## Results

### Ablation of type I interferon signaling and human FcRn expression are required for echovirus infection of the brain

Previously, we developed a mouse model of echovirus pathogenesis through intraperitoneal (IP) inoculation using mice expressing the human homologue of hFcRn (hFcRn ^Tg32^) that are also ablated in type I IFN signaling by deletion of IFNAR (hFcRn^Tg32^-IFNAR^-/-^) (21). hFcRn ^Tg32^ mice are deficient in expression of mouse FcRn and express human FcRn under the control of the native human promotor (29). To determine if type III IFNs also play a role in echovirus infections at secondary sites of infection including the brain, we generated hFcRn ^Tg32^ mice deficient in IFNLR expression, the receptor for type III IFNs (hFcRn^Tg32^-IFNLR^-/-^) (30). We used six genotypes of mice, including the hFcRn mice described above (hFcRn ^Tg32^, hFcRn^Tg32^-IFNAR^-/-^, and hFcRn^Tg32^-IFNLR^-/-^) and animals expressing murine FcRn that were immunocompetent (C57/BL6, WT) or deficient in type I or III IFN signaling (IFNAR ^-/-^ or IFNLR ^-/-^, respectively) (**Figure 1A**). Neonatal (7-day old) mice were inoculated with 10^4^ PFU of echovirus 5 (E5) by the IP route and monitored for three days post-inoculation (dpi). We observed death in 100% of hFcRn^Tg32^-IFNAR^-/-^ animals by 2 dpi (**Figure 1B**). In contrast, there were no clinical symptoms of illness in any other genotype and all animals survived until 3dpi (**Figure 1B**). There were no significant differences in mortality between male and female hFcRn^Tg32^-IFNAR^-/-^ mice (**Supplemental Figure 1A**). We next determined the level of circulating virus in the blood and brains of these animals. We detected high levels of E5 in the blood (6 of 6 mice) and brains (12 of 12 mice) of hFcRn^Tg32^-IFNAR^-/-^ mice, but no detectable virus in any other genotype (**Figure 1C, 1D**). There were no significant differences in E5 titer in the brains of male or female hFcRn^Tg32^-IFNAR^-/-^ mice (**Supplemental Figure 1B**).

**Figure 1.**
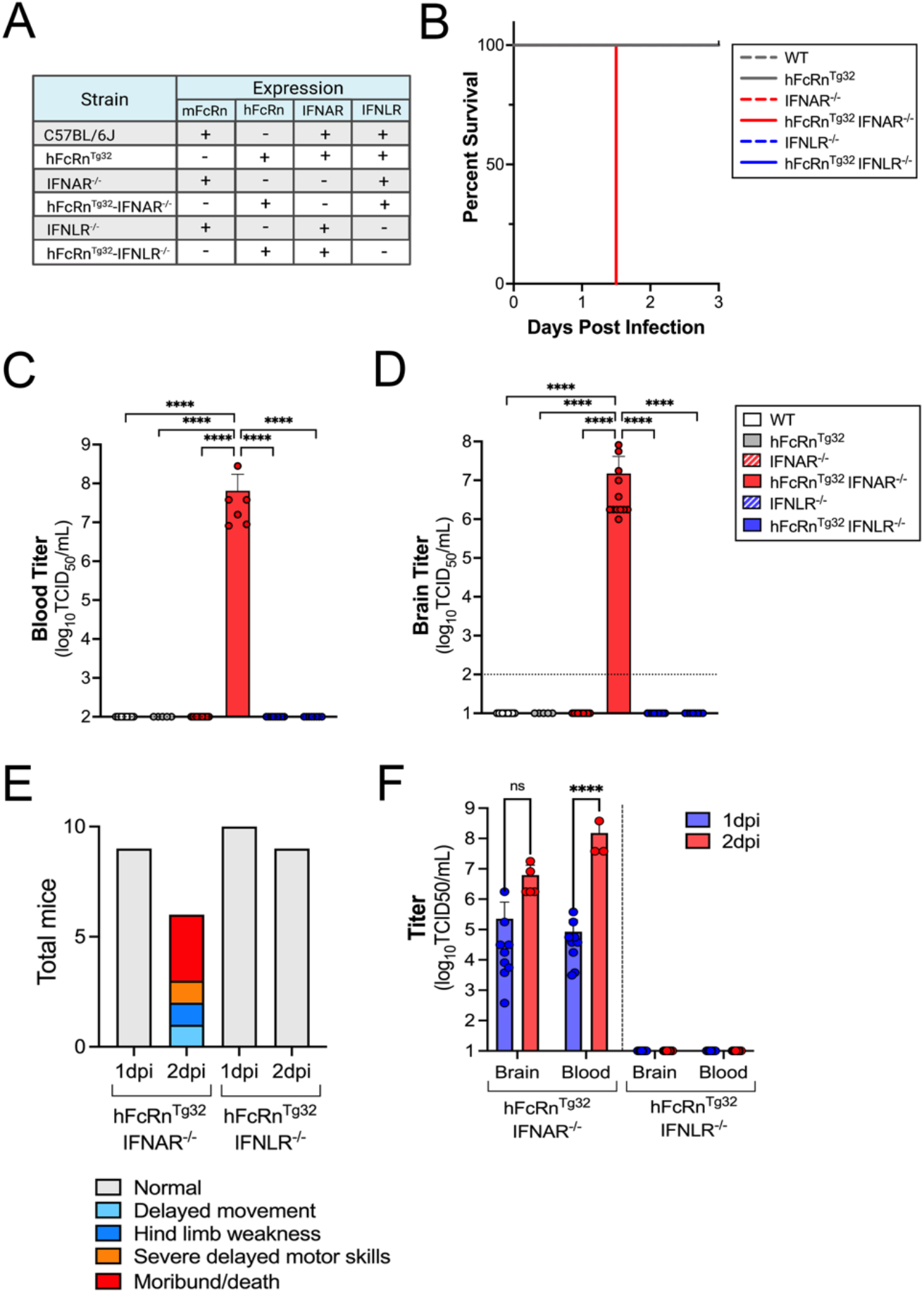
Ablation of type I interferon signaling and human FcRn expression are required for echovirus infection in the brain. **(A)** Table of the six genotypes used in this study. Shown is the expression of mouse or human FcRn, IFNAR, and IFNLR amongst these genotypes. **(B)** Survival of the indicated genotype of mice inoculated with 10^4^ PFU E5 by the IP route and monitored for 3 days post-inoculation. The number of pups in each genotype are as follows: WT (17), hFcRn ^Tg32^ (6), IFNAR ^-/-^ (17), hFcRn^Tg32^-IFNAR^-/-^ (12), IFNLR ^-/-^ (14), and hFcRn^Tg32^-IFNLR^-/-^(8). The log-rank test was used to analyze the statistical difference of the survival rate. **(C and D)**. At 3dpi, animals were sacrificed and viral titers in blood **(C)** and brain **(D)** determined by TCID50 assays. Titers are shown as log_10_TCID50/mL with the limit of detection indicated by a dotted line. Data are shown as mean ± standard deviation with individual animals shown as each data point. Data are shown with significance determined with a Kruskal-Wallis test with a Dunn’s test for multiple comparisons. **(E and F)** hFcRn^Tg32^-IFNAR^-/-^ and hFcRn^Tg32^-IFNLR^-/-^ mice were inoculated with 10^3^ PFU of E5 by the IP route and monitored for signs of disease. **(E)** Clinical symptoms observed of hFcRn^Tg32^-IFNAR^-/-^ or hFcRn^Tg32^-IFNLR^-/-^ pups at either one- or two-days post infection. **(F)** At 1 or 2dpi, animals were sacrificed and viral titers in blood and brain determined by TCID50 assays. Titers are shown as log_10_TCID50/mL with the limit of detection indicated by a dotted line. Data are shown as mean ± standard deviation with individual animals shown as each data point. Data are shown with significance determined with a Two-way Anova with Šídák’s multiple comparisons tests (*p<0.05, **p<0.005, ***p<0.0005, ****p<0.0001).

Consistent with our previous findings with E11 (21), we found that there was robust replication in the livers and pancreases of E5 infected hFcRn^Tg32^-IFNAR^-/-^ mice (12 of 12 mice) (**Supplemental Figure 1C, 1D**). However, in contrast to the brain, we found that hFcRn^Tg32^-IFNLR^-/-^ animals also contained virus in the liver (4 of 8 mice) and pancreas (8 of 8 mice) (**Supplemental Figure 1C, 1D**). We also detected low to mid-levels of E5 in the pancreases of immunocompetent hFcRn ^Tg32^ mice (5 of 5 mice) and to a much lesser extent in liver (1 of 5 mice) (**Supplemental Figure 1C, 1D**). There was no detectable virus in any animals expressing the murine homologue of FcRn (**Supplemental Figure 1C, 1D**), consistent with our previous work (21). Collectively, these data show that echovirus infections in the brain require expression of hFcRn and that the primary barrier to infection is type I IFN-mediated signaling.

### Echovirus infections cause paralysis and motor defects in infected mice

Because we observed high levels of mortality in mice infected with 10^4^ PFU E5 by the IP route, we investigated the neurotropism and neurovirulence of E5 in mice infected with a lower inoculum (10^3^ PFU) of E5 by the IP route and monitored animals daily for 2 days. At this lower inoculum we observed death in approximately 50% of hFcRn^Tg32^-IFNAR^-/-^ animals by 1.5-2 days dpi compared to no mortality in hFcRn^Tg32^-IFNLR^-/-^ animals (**Supplemental Figure 2A**). There were no differences in mortality between male and female hFcRn^Tg32^-IFNAR^-/-^ mice (**Supplemental Figure 2C**). Infected animals were monitored for signs of illness (e.g., delayed movements, paralysis, discoloration, lack of nursing, lack of parental care, and death) throughout the duration of infection. There were no signs of clinical illness in echovirus-infected hFcRn^Tg32^-IFNLR^-/-^ animals at 1 or 2 dpi (**Figure 1E**). There were also no obvious clinical symptoms in hFcRn^Tg32^-IFNAR^-/-^animals at 1dpi. However, by 1.5-2dpi, there were clear defects in motor skills and various degrees of paralysis in hFcRn^Tg32^-IFNAR^-/-^ infected animals. There were a range of defects ranging from mild loss of motor function in one or more limb characterized by difficulty walking, hemiplegia that obstructed mobility or hind limb paralysis (**Figure 1E** and **Supplemental Figure 2B**). To correlate clinical symptoms with infection, we titrated virus from the brains of infected animals at 1 and 2 dpi, which revealed similar titers at both days in brain and higher titers in blood at 2dpi in hFcRn^Tg32^-IFNAR^-/-^ animals (**Figure 1F**). There were no differences in titers between male and female mice (**Supplemental Figure 2D**). Additionally, similar to the lack of clinical disease and low morality rates, no virus was detected in the blood or brains of hFcRn^Tg32^-IFNLR^-/-^ animals (**Figure 1F**). These data show that echoviruses induce clinical symptoms of neurological disease, which occurs in an hFcRn-dependent manner and requires ablation of type I IFN signaling.

### Immunological signature of echovirus infected brains

To define the immunological signature of echovirus infected brains, we harvest the brains of E5-infected mice expressing hFcRn (hFcRn ^Tg32^, hFcRn^Tg32^-IFNAR^-/-^, and hFcRn^Tg32^-IFNLR^-/-^) at 3dpi and performed multianalyte Luminex-based profiling of 27 cytokines and chemokines on brain tissue homogenates. We found that echovirus infection induced high levels of cytokines in brain tissue of hFcRn^Tg32^-IFNAR^-/-^ infected mice at 3dpi, including high levels of Granulocyte colony-stimulating factor-3 (GSF-3), which was the most abundant cytokine detected (∼750-fold over uninfected) (**Figure 2A, 2C**). Other highly induced cytokines included IL-6 (25-fold over uninfected), CXCL10 (17-fold over uninfected), monocyte chemoattractant protein-1 (MCP-1, 16-fold over uninfected), and keratinocyte-derived chemokine (KC, 12-fold over uninfected) (**Figure 2A, 2D, 2E, 2F**). There was no significant induction of any cytokines in infected hFcRn ^Tg32^ or hFcRn^Tg32^-IFNLR^-/-^ animals at 3dpi (**Figure 2A**). We did not detect the type I IFNs IFN-α2 or IFN-β or the type III IFN IFN-λ1 in the brains of any mice (**Supplemental Figure 3A-C**).

**Figure 2.**
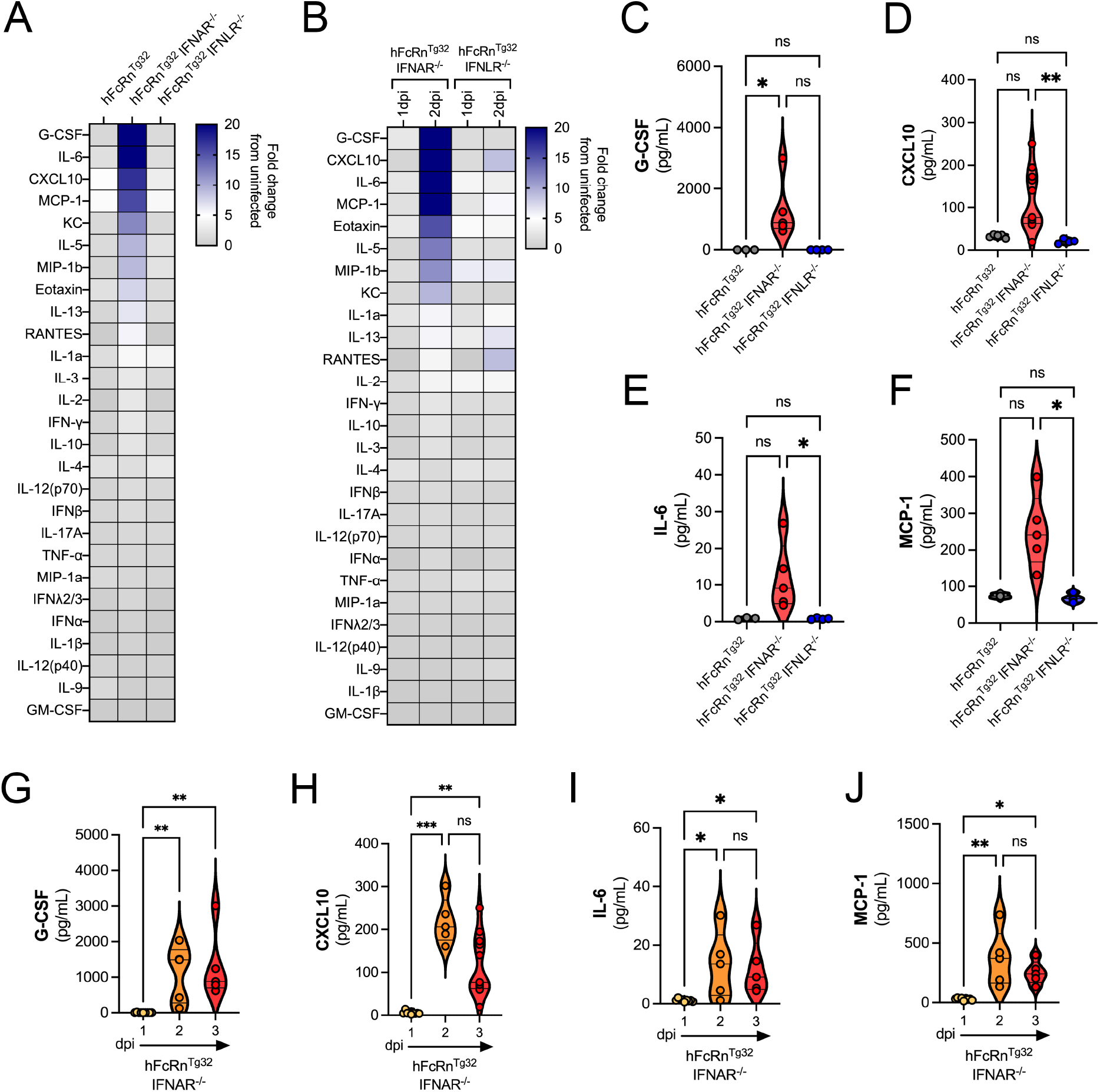
Immunological signature of echovirus infected brains. Pups were IP inoculated and sacrificed either 1, 2, or 3dpi. Cytokine expression in brain tissue homogenates were analyzed by multiplex Luminex assays. **(A)** Heatmap demonstrating the induction (shown as fold-change from uninfected control) in E5-infected mice of the indicated genotype sacrificed at 3dpi. Blue denotes significantly increased cytokines in comparison to untreated. Grey or white denote little to no changes (scale at top right). **(B)** Heatmap demonstrating the induction (shown as fold-change from hFcRn ^Tg32^ pups) in E5-infected mice of the indicated genotype sacrificed at 1 or 2dpi, as indicated. Blue denotes significantly increased cytokines in comparison to untreated. Grey or white denote little to no changes (scale at top right). **(C-F)** The top four cytokines induced from animals sacrificed at 3dpi from (A) including G-CSF **(C)**, CXCL10 **(D)**, IL-6 **(E)**, MCP-1 **(F)** shown as pg/mL. **(G-J)** The top four cytokines induced in hFcRn^Tg32^-IFNAR^-/-^ animals over the course of the infection time. Shown are G-CSF **(G)**, CXCL10 **(H)**, IL-6 **(I)**, and MCP-1 **(J)**. Data are shown as mean ± standard deviation and individual animals (points). Data are shown with significance determined with a Kruskal-Wallis test with a Dunn’s test for multiple comparisons (*p<0.05, **p<0.005, ***p<0.0005, ****p<0.0001, ns-not significant).

We next assessed the kinetics of cytokine induction in echovirus infected brains isolated from hFcRn^Tg32^-IFNAR^-/-^ and hFcRn^Tg32^-IFNLR^-/-^ infected animals at 1 and 2dpi. Consistent with the overall low levels of virus in the brains of mice at 1dpi, we observed very little cytokine induction in hFcRn^Tg32^-IFNAR^-/-^ animals at this time (**Figure 2B**). However, by 2dpi, there were high levels of similar cytokines observed at 3dpi, including G-CSF (647-fold over uninfected), CXCL10 (33-fold over uninfected), IL-6 (28-fold over uninfected), and MCP-1 (23-fold over uninfected) (**Figure 2B, 2G-J**). This cytokine induction paralleled the accumulation of viral RNA (vRNA) in the brain, as assessed by RT-qPCR for vRNA and the transcript for CXCL10 (**Supplemental Figure 3D-E**). There were no cytokines significantly induced in infected hFcRn^Tg32^-IFNLR^-/-^ animals at either time point (**Figure 2B**). These data show that echovirus infection of brain tissue results in an immunological response characterized by the induction of select pro-inflammatory cytokines and chemokines.

### Echoviruses replicate in the leptomeninges to induce meningeal inflammation

Very little is known regarding the cellular or structural targets of echovirus infections in the brain, including the meninges, which is composed of three membranous layers. To define the site(s) of echovirus replication in the brain, we infected hFcRn ^Tg32^, hFcRn^Tg32^-IFNAR^-/-^, and hFcRn^Tg32^-IFNLR^-/-^ mice with 10^4^ PFU E5 for 3d to maximize infection. At this time, whole brains were removed, sectioned, and processed for hybridization chain reaction (HCR) using E5-specific probes. HCR allows for fluorescent quantitative RNA detection with enhanced sensitivity over conventional hybridization approaches given signal amplification resulting from the self-assembly of secondary detection hairpins into amplification polymers (31, 32). Whole brain confocal microscopy-based tile scanning of ∼36mm ^2^ was then performed to define the region(s) and cell(s) infected by E5. We observed high levels of E5 vRNA in infected hFcRn^Tg32^-IFNAR^-/-^ brain tissue, but none in hFcRn ^Tg32^ or hFcRn^Tg32^-IFNLR^-/-^ brains, at 3dpi, which was localized to a distinct region surrounding the brain (**Figure 3A, Supplemental Figure 4A, 4B**). This region was specific for the leptomeninges, which includes the two inner layers of the meninges (**Figure 3B, 3C**). In addition, there were concentrated regions of high levels of E5 vRNA surrounding blood vessels localized throughout the meninges (**Figure 3D**). Although the choroid plexus has been reported to express high levels of FcRn (15), we did not detect any infection in this region (**Supplemental Figure 4C)**. To determine the timing of E5 vRNA in the meninges, we performed HCR in whole brains isolated from hFcRn^Tg32^-IFNAR^-/-^ mice infected with E5 for 1-3dpi. At 1dpi, we found that the levels of E5 vRNA were undetectable like uninfected controls, but by 2dpi, there were clear areas of E5 infection, which significantly increased by 3dpi (**Figure 3E, 3F**).

**Figure 3.**
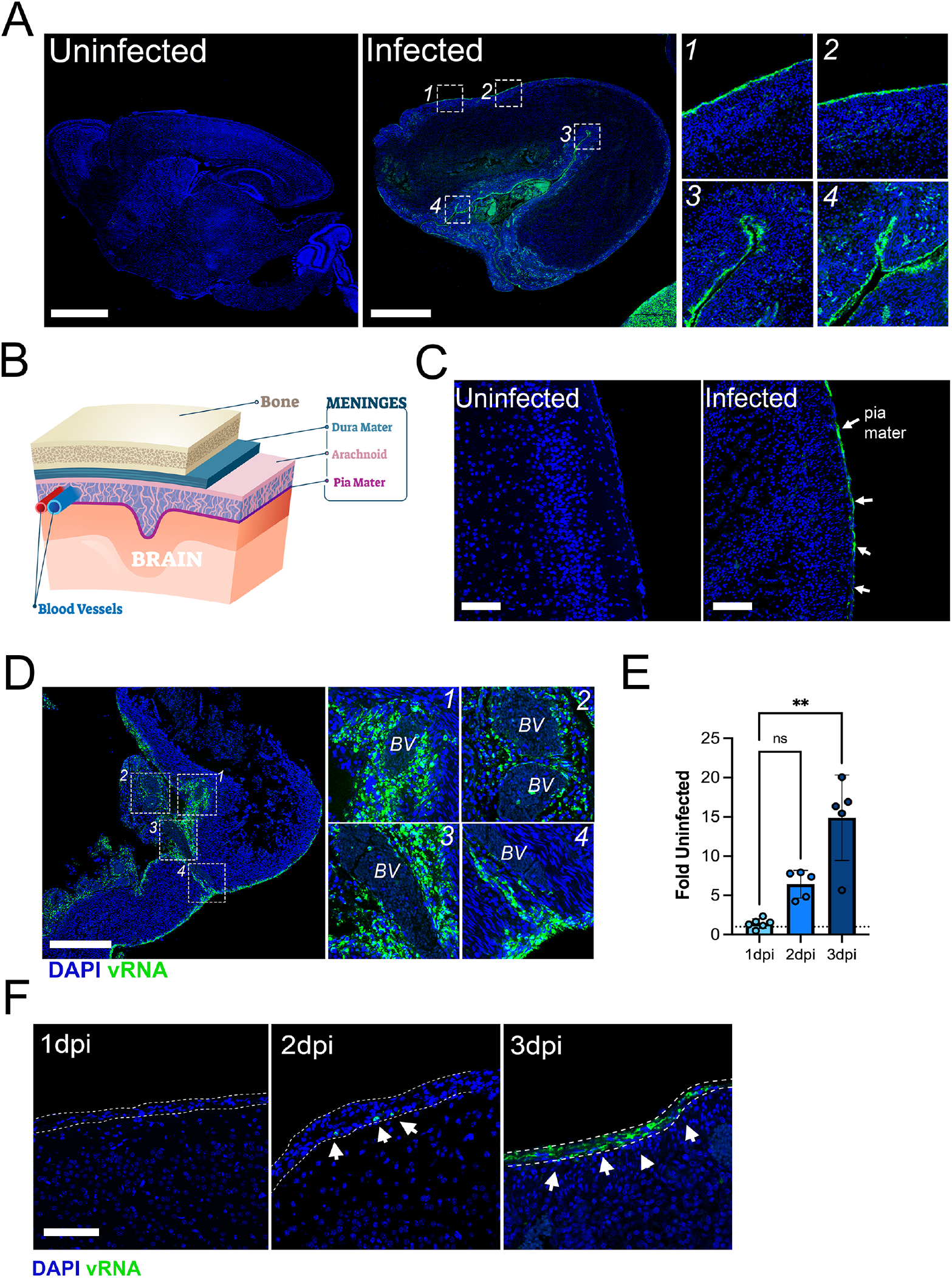
Echovirus replication in the leptomeninges. **(A)** Tile scan of the brains from an uninfected animal (left) and E5 infected hFcRn^Tg32^-IFNAR^-/-^ animal at 3dpi (middle) using Hybridization chain reaction RNA-FISH (HCR) for vRNA (in green) and DAPI (in blue). Numbered white boxes show zoomed areas to the right. **(B)** Schematic representation of the different layers of the meninges surrounding the brain. The dura mater in teal with the leptomeninges (arachnoid and pia mater) in shades of pink. **(C)** HCR for vRNA (in green) and DAPI (in blue) of a brain from an uninfected or an E5 infected hFcRn^Tg32^-IFNAR^-/-^ animal at 3dpi. White arrows denote areas of vRNA. **(D)** Tile scan of HCR for E5 vRNA from a brain of an infected hFcRn^Tg32^-IFNAR^-/-^ animal at 3dpi of areas of infection surrounding blood vessels (BV). DAPI-stained nuclei in blue and vRNA in green. White boxes indicate zoomed images at right with numbers in the top right corner denoting the corresponding zoomed image. **(E)** Image analysis of the extent of vRNA signal in the brains of hFcRn^Tg32^-IFNAR^-/-^ animals at 1-3dpi as shown as a fold change from uninfected controls. Symbols represent unique regions used in quantification. Data are shown with significance determined with a Kruskal-Wallis test with a Dunn’s test for multiple comparisons (**p<0.01, ns-not significant). **(F)** HCR for vRNA (in green) and DAPI (in blue) of the brains from infected hFcRn^Tg32^-IFNAR^-/-^ animals at 1, 2, or 3dpi, as indicated at top left. Dotted lines highlight the leptomeninges and arrows denote infected cells. Scale bars are as follows: 1mm (A), 100μm (C), 1mm (D), and 50μm (F).

To define the localization of FcRn within the brains of hFcRn ^Tg32^ mice and to correlate this expression with E5 infection, we performed immunohistochemistry for hFcRn in hFcRn^Tg32^-IFNAR^-/-^ mice. We found that hFcRn was enriched in the leptomeninges of infected animals, where it was concentrated to regions associated with high levels of vRNA (**Figure 4A**).

**Figure 4.**
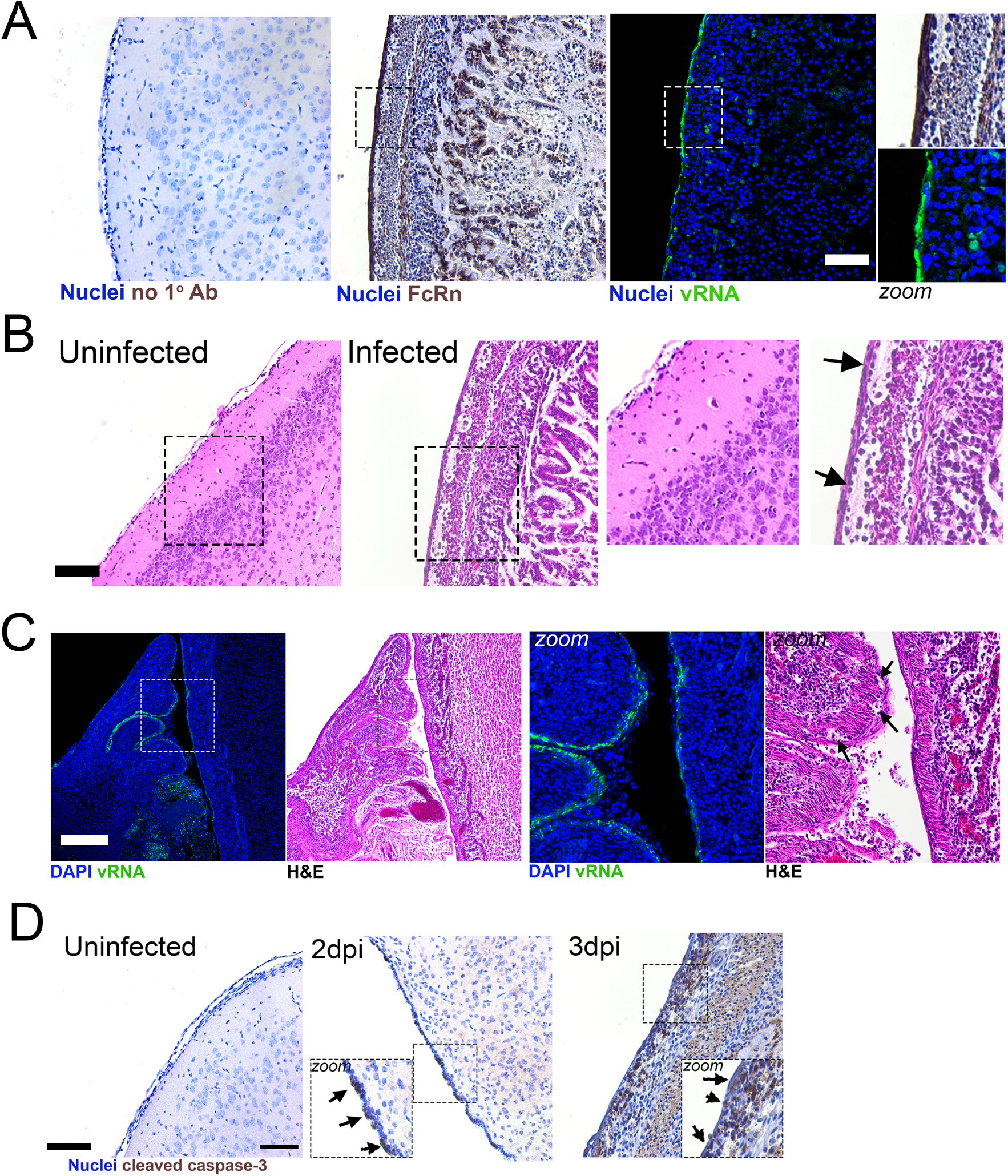
Echovirus replication in the meninges induces inflammation and cell death. **(A)**, Immunohistochemistry for hFcRn and HCR for vRNA in the brain of a hFcRn^Tg32^-IFNAR^-/-^ animal infected with E5 for 3d. At left, no primary antibody control in an uninfected control animal. Middle, hFcRn IHC and right, HCR for vRNA (in green) and DAPI (in blue) from the same E5 infected hFcRn^Tg32^-IFNAR^-/-^ animal. Boxes indicate regions that are zoomed at right. **(B)** Hematoxylin and eosin staining of representative brain sections from an uninfected or E5 infected hFcRn ^Tg32^- IFNAR ^-/-^ animal. Black boxes show areas of zoomed images at right. Arrows denote regions of inflammation within the meninges. **(C)** HCR for E5 vRNA or hematoxylin and eosin staining of the cerebellum within the brain of a hFcRn^Tg32^-IFNAR^-/-^ animal at 3dpi. HCR shows vRNA (in green) and DAPI (in blue). Boxes denote areas of zoomed images at right. Arrows denote region of inflammation within the meninges. **(D)** IHC for cleaved caspase 3 in uninfected, or hFcRn^Tg32^-IFNAR^-/-^ animals infected with E5 for 2dpi, or 3dpi. Arrows denote cells that are positive for cleaved caspase 3. Black boxes show zoomed images. Scale bars are as follows: 100μm (A, B, D) and 1mm (C).

Next, we assessed the impacts of E5 infection on the integrity of the meninges. We noted instances of acute meningitis and inflammatory tissue damage in hFcRn^Tg32^-IFNAR^-/-^ mice infected with E5 for 3dpi, which included areas of immune cell infiltration (**Figure 4B**). Areas of inflammation correlated with high levels of E5 replication, as indicated by aligning HCR and H&E images (**Figure 4C**). Lastly, to determine whether E5 infection induces direct cell death or damage to the meninges, we performed immunohistochemistry for cleaved caspase-3, which revealed discrete areas of apoptosis as early as 2dpi, with more significant levels at 3dpi (**Figure 4D**). Together, these data show that echovirus infection of the meninges induces pronounced tissue damage and inflammation.

## Discussion

Echovirus infections are common causes of neonatal meningitis, which can be fatal. However, how these viruses infect the brain and their regional tropism within the brain remains unknown. Here, we developed an *in vivo* model of echovirus-induced meningitis and used this model to define the cellular targets and host immune pathways associated with echovirus infection within the brain. We show that expression of the human homologue of FcRn is necessary, but not sufficient, for echovirus infection of the brain. In addition, we show that type I IFNs, but not type III IFNs, provide a barrier to echovirus infections of the meninges and that ablation of this pathway sensitizes the brain to infection, which induces a robust immune response. Lastly, we define the specificity of echovirus replication in the brain and show that high levels of replication in the leptomeninges induced inflammation and cell death. These studies thus provide key insights into the events associated with echovirus-induced meningitis and an *in vivo* model that could be used to test echovirus therapeutics targeting echovirus-induced neuronal disease.

FcRn is expressed on the microvasculature of the blood-brain barrier (15), where it has been proposed to mediate the efflux of antibodies out of the brain (33). In addition to this localization, FcRn is expressed in the epithelium of the choroid plexus (15). This pattern of expression is conserved between humans and in hFcRn ^Tg32^ mice (34). In addition to these sites, our data show that FcRn is also expressed at distinct regions of the meninges. The meninges are composed of diverse cell types, with distinct patterns of expression regionally between the dura, arachnoid layer, and pia mater, with the arachnoid and pia mater referred to as the leptomeninges (35). The meningeal layers are primarily comprised of fibroblasts, with each subtype expressing their own unique transcriptional signature (35). In addition to fibroblasts, the arachnoid layer also contains arachnoid barrier cells, which are epithelial-like in origin and separate the dura mater from the subarachnoid space in which CSF accesses the brain. The meninges also contain a large population of immune cells, including CD4 ^+^ T cells and B cells. Notably, meningeal macrophages are a specialized subclass of macrophages which are long-lived in the leptomeninges (36). We observed the highest levels of E5 infection restricted to the leptomeninges, the inner layer of the meninges containing the pia mater and arachnoid layers, which also expressed high levels of FcRn. Given the robust levels of E5 infection in the meninges, it is likely that fibroblasts are at least one target of infection. However, it is possible that other cell types, such as macrophages, are also permissive to infection and contribute to pathogenesis. Collectively, these data suggest that the regional expression of FcRn plays a direct role in the tropism of echoviruses in the brain and directly correlates with their high levels of replication in the meninges.

Several studies have investigated the levels of cytokines and chemokines within the CSF of infants and children with confirmed echovirus-induced meningitis. The CSF of children with echovirus 30-induced meningitis contain very high levels of MCP-1 and IL-6 compared to controls (23, 37). Our data show that hFcRn^Tg32^-IFNAR^-/-^ animals also induce very high levels of MCP-1 and IL-6 in brain tissue homogenates. MCP-1, also known as CCL2, is a monocyte chemoattractant responsible for recruiting monocytes and dendritic cells to the site of inflammation due to infection. IL-6 is a proinflammatory cytokine that is secreted by macrophages that have detected a viral infection. These immune mediators play important roles in alerting the immune system to a viral infection and recruiting immune cells to the sites of infection. However, because we used brain tissue homogenates and human studies can are restricted to CSF, it is unclear which cytokines induced by echovirus infections mediate the influx of immune cells observed in echovirus-induced meningitis. Although the differential roles of type I and III IFNs has been established in the respiratory and intestinal epithelium during enterovirus infection (30, 38-40), whether these IFNs play distinct roles in enterovirus-induced meningitis has remained unclear. Our data suggest that type I IFNs are the sole barrier to echovirus infection of the CNS, which is consistent with previous work defining indicating that these IFNs form at least one bottleneck to poliovirus access to the CNS (41). However, type I IFNs were not detected in multiplex Luminex profiling of brain homogenates from echovirus-infected mice, suggesting that these IFNs are not produced locally in the brain. It is therefore likely that circulating IFNs produced at distal sites, such as the liver, trigger antiviral responses in the CNS that protect from echovirus infections.

In humans, neonates are at increased risks for echovirus-induced morbidity and mortality. For example, rates of echovirus-induced paralysis decreased with increased age at inoculation (42). Our previous work showed that adult echovirus-infected mice exhibit low levels of infection in the brain following IP inoculation (21). Our data presented here show that neonatal mice are highly sensitive to echovirus-induced neuronal dysfunction and contain high levels of viral infection within the meninges. The mechanistic basis for age-related differences in echovirus infection of the brain remains unclear. Given that BBB forms in mice prior to birth (43), this barrier is unlikely to participate in these age-related differences. Age-related differences in neuronal susceptibility to infections not specific to echoviruses and has also been shown for reovirus (44). However, unlike this work, our data suggest that type I IFNs are not the primary drivers for differential age susceptibility given that both adult and neonatal animals in our work were deficient in IFNAR expression. Other work has shown that age-related differences in susceptibility to cardiac infections by CVB may result from differences in receptor expression (45, 46). Age-related differences in FcRn expression in humans are unknown, but its expression in human liver is not dependent on age (47), suggesting this is unlikely to explain age-related differences in echovirus-associated neuronal infections. Therefore, it remains unclear which factors contribute to age-related difference in infection of the CNS.

Our findings presented here define key aspects of echovirus infection of the brain, including the sites of viral replication and the consequences of this infection. We show that FcRn is necessary but not sufficient for echovirus infection of the brain *in vivo* and that type I IFNs control infection within the meninges. Collectively, these studies develop a model of echovirus aseptic meningitis, which could aid in the testing of novel therapeutics.

## Materials and Methods

### Cell lines and viruses

HeLa cells (clone 7B) were provided by Jeffrey Bergelson, Children’s Hospital of Philadelphia, Philadelphia, PA, and cultured in MEM supplemented with 5% FBS, non-essential amino acids, and penicillin/streptomycin. Experiments were performed with echovirus 5 (Noyce strain, E5), which was obtained from the ATCC. Virus was propagated in HeLa cells and purified by ultracentrifugation over a 30% sucrose cushion, as described previously (48). All viruses were sequenced for viral stock purity following propagation. Purity of all viral stocks was confirmed by Sanger sequencing of VP1 using enterovirus-specific primers (49). Briefly, RNA extraction was performed on 10μl of purified virus stock, according to manufacturer’s instructions (Qiagen Cat. 529904). RNA was reverse transcribed using SuperScript III reverse transcription kit, (Invitrogen cat. 18080093) according to manufacturer’s instructions, with a pan enterovirus primer (vir21; ATAAGAATGCGGCCGCTTTTTTTTTTTTTTTTTTTTTTTTT), followed by an RNaseH treatment for 20 minutes at 37°C. PCR was performed with 5μl of the cDNA reaction using BioRad iTaq DNA polymerase (BioRad cat. 1708870). Virus specific primers are as follows: E5 forward 5’-TATCGCCAATTACAACGCGAA-3’; E5 reverse 5’-TTGGTTTGAAGTAAACCCTTA-3’.

### Animals

All animal experiments were approved by the Duke University Animal Care and Use Committees, and all methods were performed in accordance with the relevant guidelines and regulations. C57BL/6J (WT, cat. no. 000664), B6.Cg-*Fcgr* ^*t*tm1Dcr^Tg(FCGRT)32Dcr/DcrJ (hFcRn ^Tg32^, cat. no. 014565), and B6.(Cg)-Ifnar1 ^tm1.2Ees^/J (IFNAR ^-/-^, cat. no. 028288) mice were purchased from The Jackson Laboratory. hFcRn^Tg32^-IFNAR^-/-^ mice were generated as described previously (21). B6.Ifnlr ^-/-^/J (IFNLR ^-/-^) mice were provided by Dr. Megan Baldridge (Washington University School of Medicine). hFcRn^Tg32^-IFNLR^-/-^ mice were generated by crossing B6.Cg-*Fcgr* ^*t*tm1Dcr^Tg(FCGRT)32Dcr/DcrJ (hFcRn ^Tg32^, cat. no. 014565) mice with B6.Ifnlr ^-/-^/J mice (30). Breeders were established that were deficient in mouse FcRn and IFNLR and were homozygous for the hFcRn transgene. All animals used in this study were genotyped by Transnetyx.

### Suckling pup infections

7-day-old mice were inoculated by the IP route with 10^4^ or 10^3^ PFU of E5. Inoculation was performed using a 1mL disposable syringe and a 27-gauge needle in 50μL of 1X PBS. Mice were euthanized at either 1-, 2-, or 3-days post inoculation and organs harvested into 0.5mL of DMEM and stored at -80°C. Tissue samples for viral titration were thawed and homogenized with a TissueLyser LT (Qiagen) for 5 minutes, followed by brief centrifugation for 5 minutes at 8000xg. Viral titers in organ homogenates were determined by TCID50s in HeLa cells and enumerated following crystal violet staining.

### Luminex assays

Luminex profiling was performed on whole brain tissue homogenates where were homogenized with a TissueLyser LT (Qiagen) for 5 minutes, followed by a centrifugation of 10,000xg for 10 minutes. Luminex kits that were used are as follows a custom mouse IFN kit (IFN alpha, IFN beta, IL-28, Invitrogen), mouse cytokine 23-plex (Bio-Rad, M60009RDPD), and mouse CXCL10 (Invitrogen, EPX01A-26018-901), according to the manufacturer’s protocol. Assays were read on a BioPlex 200 by BioRad. Heat maps were generated using the fold change in concentration (picograms/milliliter) of each animal compared to the average of uninfected animals and was made in GraphPad Prism. Violin plots are shown as the concentration for each animal (one point) in picograms/milliliter.

### RNA Extraction and RT-qPCR

Total RNA was isolated from brains using the Sigma GenElute Total Mammalian RNA Miniprep Kit (Sigma, RTN350), according to the manufacturer protocol with the addition of a Sigma DNase digest reagent (Sigma, DNASE70). RNA (1 ug total) was reverse transcribed using iScript cDNA Synthesis Kit (Bio-Rad, 1708891) and diluted to 100 ul in ddH_2_O for subsequent qPCR analyses. RT-qPCR was performed using the iTaq Universal SYBR Green Supermix (Bio-Rad, 1725121) on a CFX96 Touch Real-Time PCR Detection System (Bio-Rad, 1855195). Primer sequences can be found in Supplemental Table 1.

### Immunohistochemistry

Tissues were fixed in 10% buffered formalin for 24hrs and then transferred to 70% ethanol. Tissues were embedded in paraffin and sectioned. Slides were stained with cleaved caspase 3 (Asp175) (9661, Cell Signaling) or human FCGRT (Abcam, ab139152). Tissue sections were deparaffinized with xylene and rehydrated with decreasing concentrations of ethanol (100%, 95%, 80%), then washed with ddH_2_O. Antigen unmasking was performed with slides submerged in 10 mM citrate buffer (pH 6.0) and heated in a steamer for 20 minutes at ∼90°C. Slides were cooled to room temperature and slides were immunostained with cleaved caspase 3 or FCGRT using Vectastain Elite ABC HRP (Vector Biolabs, PK-6100), according to the manufacturer’s instructions. Slides were incubated in 6% H_2_O _2_ in methanol for 30 min then washed 3 times for 5 minutes in H_2_O. Avidin block (Vector, SP-2001) was applied for 15 minutes and washed twice in H_2_O followed by biotin block (Abcam, ab156024) for 15 minutes and washed twice in H_2_O. Finally, serum-free protein block was applied for 10 minutes, and cleaved caspase 3 antibody was diluted 1:100 and the FCGRT antibody was diluted to 1:200 in TBS-T (Tris-buffered saline, 0.1% Tween 20) and slides incubated overnight in a humidified chamber at 4C. Next, slides were washed three times for 5 min in PBST and exposed to the goat anti-rabbit biotinylated secondary antibody (Vector, BA-1000) for 30 min. Slides were rinsed in PBST three times for 5 min and the Vectastain Elite ABC HRP kit was applied for 30 min. Slides were rinsed in PBST for three times for 5 min and diaminobenzidine substrate for 5 mins, which was terminated with water incubation. Slides were counterstained with hematoxylin for 1 min, thoroughly rinsed with H_2_O, and incubated in 0.1% sodium bicarbonate in H_2_O for 5 mins. Slides were then dehydrated with increasing concentrations of ethanol, cleared with xylene and mounted with Cytoseal 60 (Thermo Scientific, 83104). Images were captured on an IX83 inverted microscope (Olympus) using a UC90 color CCD camera (Olympus).

### HCR and Imaging

HCR was performed following the Molecular Instruments HCR v3.0 protocol for FFPE human tissue sections (31, 32). Briefly, tissue sections were deparaffinized with xylene and rehydrated with decreasing concentrations of ethanol (100%, 95%, 80%). Antigen unmasking was performed with slides submerged in 10 mM citrate buffer (pH 6.0) and heated in a steamer for 20 minutes at ∼90°C. Slides were cooled to room temperature. Sections were treated with 10 µg/mL Proteinase K for 10 min at 37°C and washed with RNase free water. Samples were incubated for 10 minutes at 37°C in hybridization buffer. Sections were incubated overnight in a humidified chamber at 37°C with 3 pmol of initiator probes in hybridization buffer. We designed probes for E5 (**Supplemental Table 2**). The next day, slides were washed in probe wash buffer and 5x SSCT for 4x 15 min, according to the manufacturer’s instructions. Samples were incubated in a humidified chamber at 37°C for 30 minutes in amplification buffer. Fluorescent hair pins were heated to 95 °C for 90 seconds and snap cooled at room temperature for 30 min. Hairpins and amplification buffer were added to the sample and incubated overnight at room temperature. Hairpins were washed off with 5x SSCT for 5 minutes, 15 minutes, 15 minutes, and 5 minutes followed by a wash with PBS containing DAPI. Slides were mounted in Vectashield with DAPI. Slides were imaged on a Zeiss 880 with Airyscan inverted confocal microscope. Image analysis was performed using FIJI.

### Statistics

All statistical analysis was performed using GraphPad Prism version 9. Data are presented as mean ± SD. Parametric tests were applied when data were distributed normally based on D’Agostino–Pearson analyses; otherwise, nonparametric tests were applied. The log-rank test was used to analyze the statistical difference of the survival rate in kaplan meier curves. In most cases, a Kruskal-Wallis test with a Dunn’s test for multiple comparisons or Two-way Anova with Šídák’s multiple comparisons tests were used to determine statistical significance, as described in the figure legends. P values of <0.05 were considered statistically significant, *p<0.05, **p<0.005, ***p<0.0005, ***p<0.0001, as noted in figure legends.

## Acknowledgements

We thank Cristian Ovies and Kalena Grimes (Duke University School of Medicine) for technical assistance, Megan Baldridge (Washington University) for providing IFNLR ^-/-^ mice, and Sujan Shresta (La Jolla Institute for Immunology) for providing hFcRn^Tg32^-IFNAR^-/-^ mice. This project was supported by NIH R01-AI150151 (C.B.C), NIH T32-AI060525 (A.I.W), NIH F31-AI149866 (A.I.W). The funders had no role in study design, data collection and analysis, decision to publish, or preparation of the manuscript.

---

**Supplemental Figure 1.**
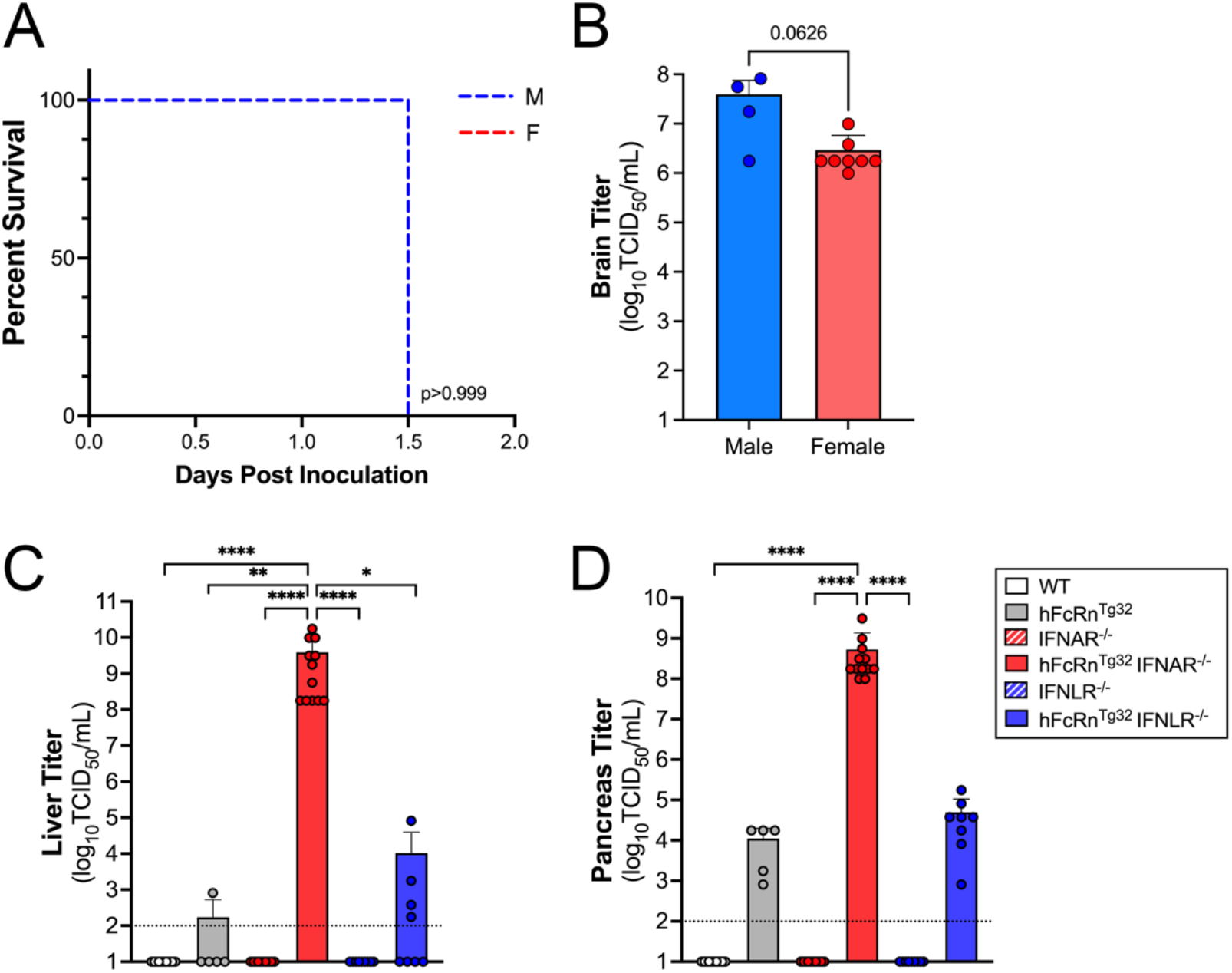
**(A)** Survival of hFcRn^Tg32^-IFNAR^-/-^ mice inoculated with 10^4^ PFU of E5 by the IP route and monitored for 3 days post-inoculation. Animals are broken down by sex (M-male, F-female) to represent any potential sex differences in mortality. The log-rank test was used to analyze the statistical difference of the survival rate with p value shown. **(B)** Brain titers from hFcRn^Tg32^-IFNAR^-/-^ animals broken down by sex. Data are shown with significance determined with a Mann-Whitney U test with p value shown. **(C and D)** The indicated genotype of mice inoculated with 10^4^ E5 by the IP route. At 3dpi, animals were sacrificed and viral titers in liver **(C)** and pancreas **(D)** determined by TCID50 assays. Titers are shown as log_10_TCID50/mL with the limit of detection indicated by a dotted line. Data are shown as mean ± standard deviation with individual animals shown as each data point. Data are shown with significance determined with a Kruskal-Wallis test with a Dunn’s test for multiple comparisons (*p<0.05, **p<0.005, ***p<0.0005, ***p<0.0001).

**Supplemental Figure 2.**
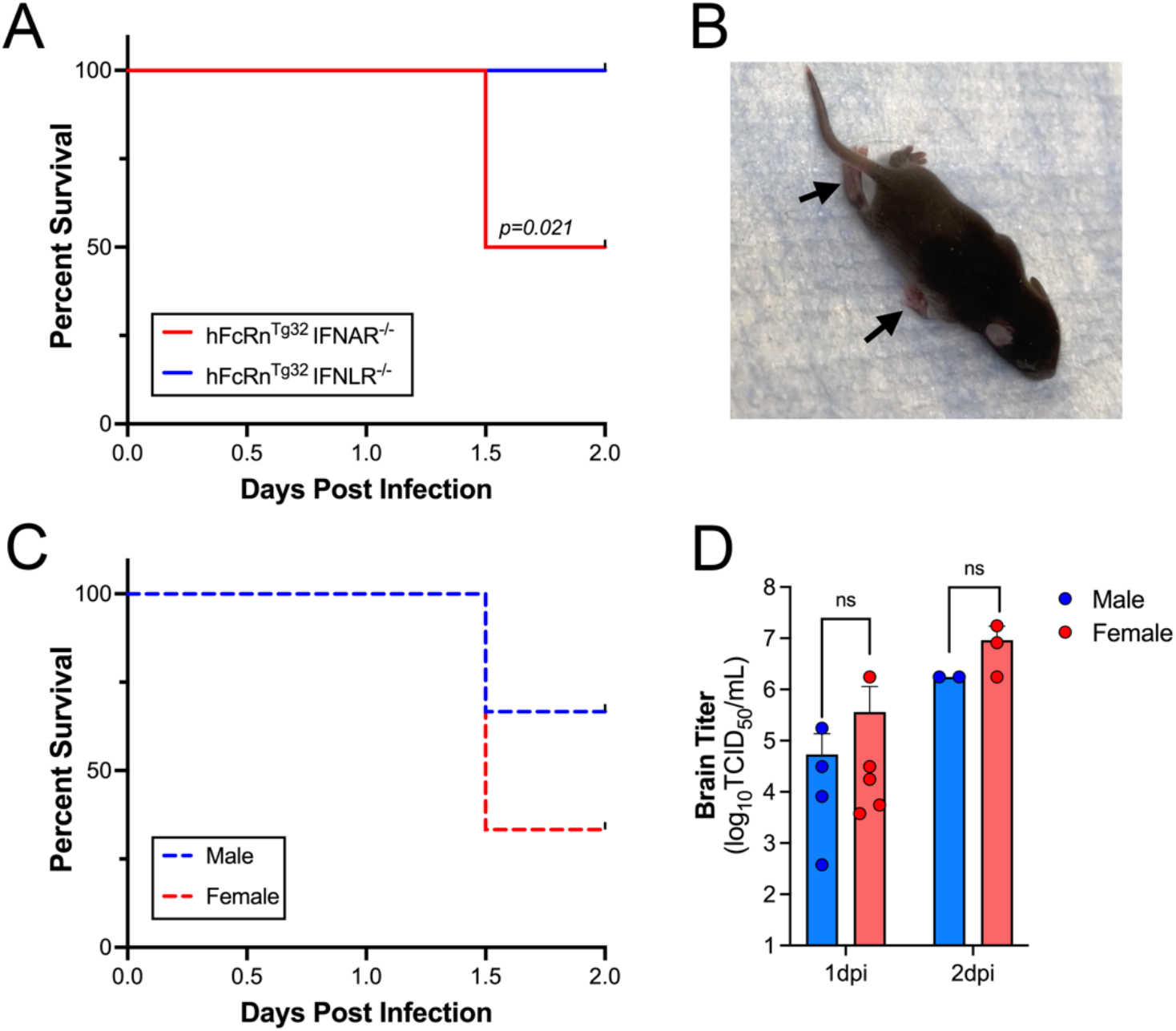
**(A)** Survival of hFcRn^Tg32^-IFNAR^-/-^ and hFcRn^Tg32^-IFNLR^-/-^ mice which were inoculated with 10^3^ PFU of E5 by the IP route and monitored for 2 days post-inoculation. The number of pups in each genotype are as follows: hFcRn^Tg32^-IFNAR^-/-^ (6) and hFcRn^Tg32^-IFNLR^-/-^ (9). The log-rank test was used to analyze the statistical difference of the survival rate of hFcRn^Tg32^-IFNAR^-/-^ or hFcRn^Tg32^-IFNLR^-/-^ pups. **(B)** Representative image of a hFcRn^Tg32^-IFNAR^-/-^ mouse with hemiplegia. Arrows denote limbs that are impacted. **(C)** Survival of hFcRn^Tg32^-IFNAR^-/-^ mice broken down by sex which were inoculated with 10^3^ PFU of E5 by the IP route and monitored for 2 days post-inoculation. The log-rank test was used to analyze the statistical difference of the survival rate of between males and females with the p value shown. **(D)** Brain titers from hFcRn^Tg32^-IFNAR^-/-^ animals broken down by sex at either 1 or 2dpi. Data are shown with significance determined with a Mann-Whitney U test, ns-not significant.

**Supplemental Figure 3.**
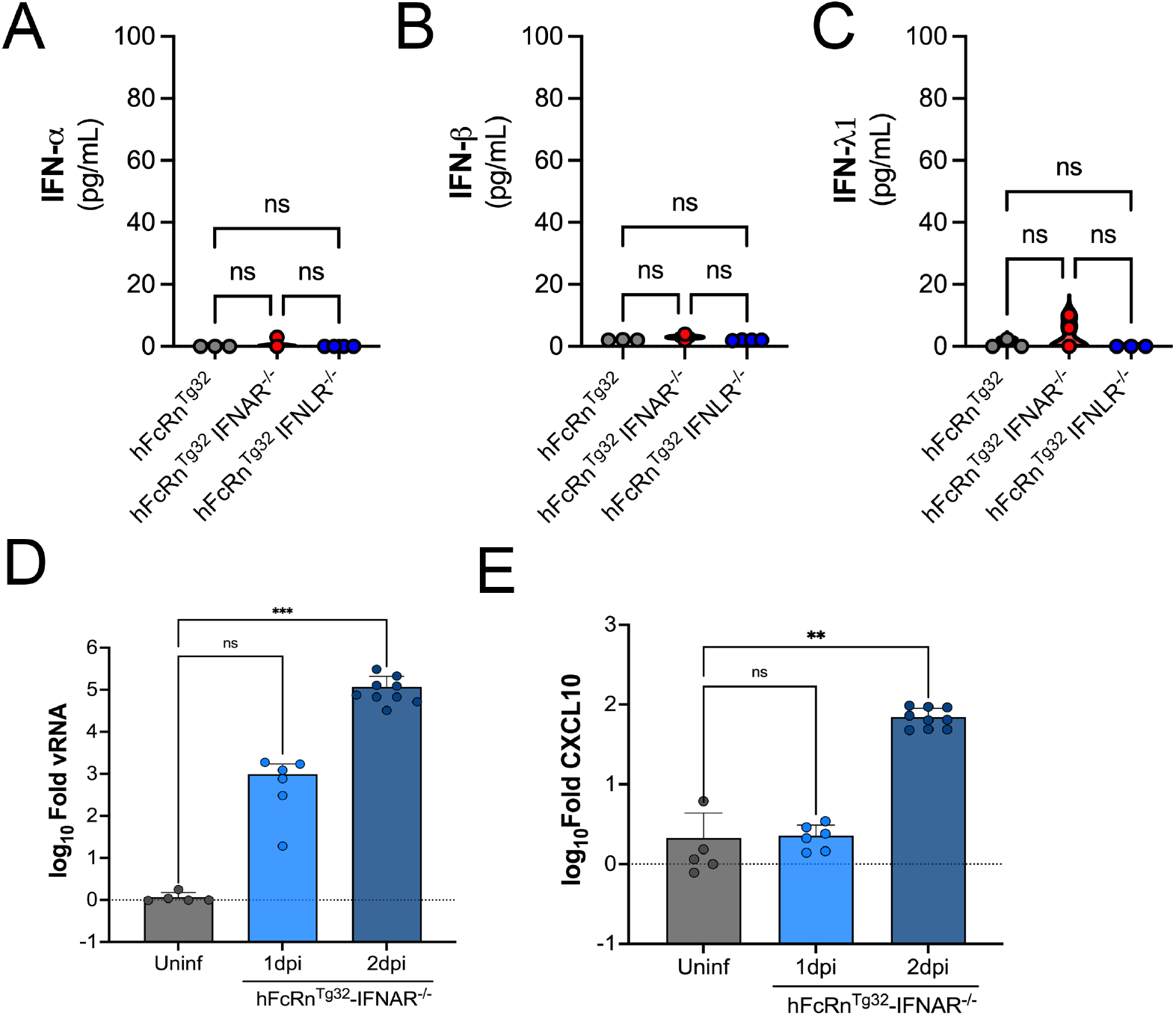
**(A-C)** Luminex multiplex assays from brain tissue homogenates of hFcRn ^Tg32^, hFcRn^Tg32^-IFNAR^-/-^, and hFcRn^Tg32^-IFNLR^-/-^ animals that were IP inoculated and sacrificed at 3dpi. Shown as the concentration in pg/mL IFN-α **(A)**, IFN-β **(B)**, and IFNλ1 **(C)**. RT-qPCR for viral RNA **(D)** or CXCL10 **(E)** from the brains of uninfected, 1dpi, or 2dpi hFcRn^Tg32^-IFNAR^-/-^ animals. Data are shown with the significance determined with a Kruskal-Wallis test with a Dunn’s test for multiple comparisons (**p<0.005, ***p<0.0005, ns-not significant).

**Supplemental Figure 4.**
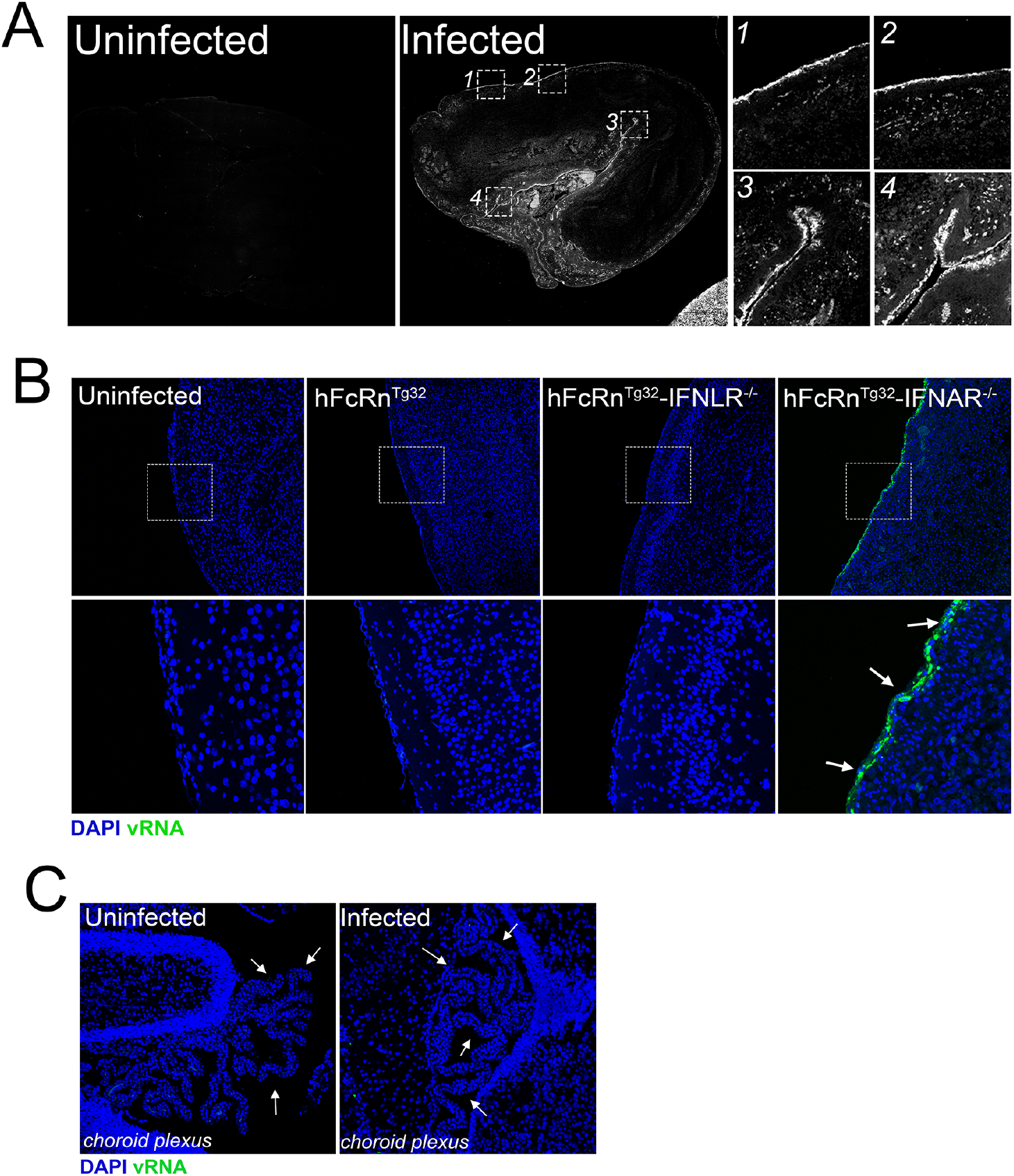
**(A)** Tile scan of the brains from an uninfected (left) and E5 inoculated at 3dpi (middle) hFcRn^Tg32^-IFNAR^-/-^ animal using Hybridization chain reaction RNA-FISH (HCR) from figure 4A. vRNA is shown in white. **(B)** HCR of brain sections from hFcRn ^Tg32^, hFcRn^Tg32^-IFNAR^-/-^, and hFcRn^Tg32^-IFNLR^-/-^ animals at 3dpi. E5 viral RNA (vRNA) is shown in green and DAPI-stained nuclei are shown in blue. White box is shown as a zoomed image below. White arrows at right denote areas of E5 vRNA. **(C)** Choroid plexus region within the brain from an uninfected (left) and E5 inoculated at 3dpi (right) hFcRn^Tg32^-IFNAR^-/-^ animal using HCR with vRNA (in green) and DAPI (in blue). White arrows show the choroid plexus.

**Supplemental Table 1.**
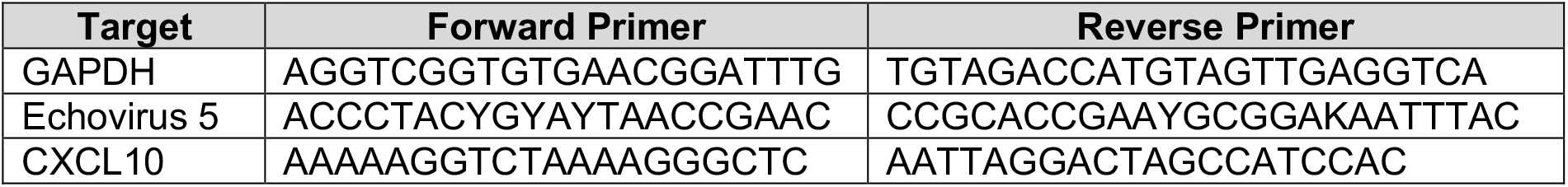
RT-qPCR Primers used in this study.

**Supplemental Table 2.**
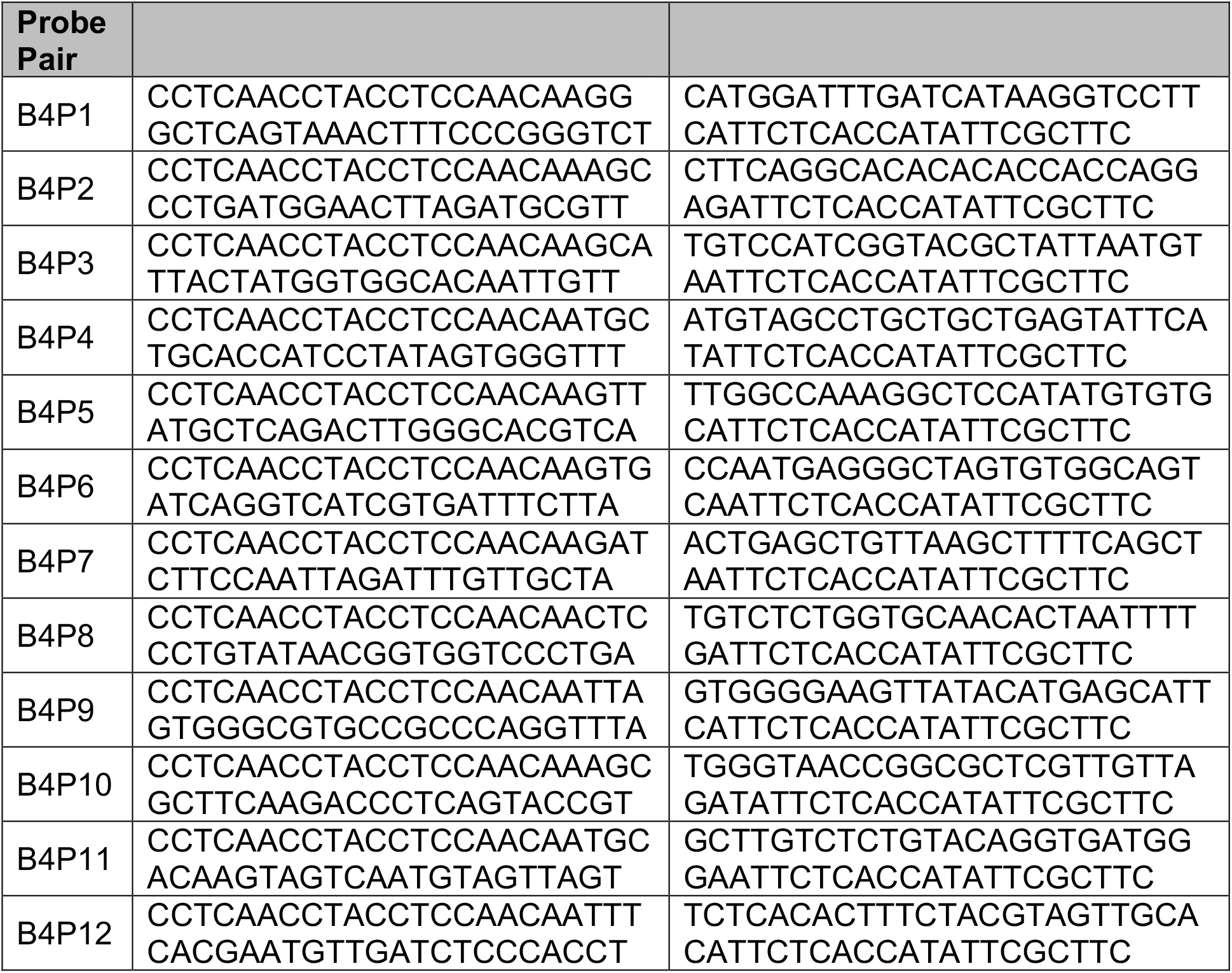
Echovirus 5 HCR Probes.

